# Smart self-propelled particles: a framework to investigate the cognitive bases of movement

**DOI:** 10.1101/2023.03.07.531552

**Authors:** Valentin Lecheval, Richard P. Mann

## Abstract

Decision-making and movement of single animals or group of animals are often treated and investigated as separate processes. However, many decisions are taken while moving in a given space. In other words, both processes are optimised at the same time and optimal decision-making processes are only understood in the light of movement constraints. To fully understand the rational of decisions embedded in an environment (and therefore the underlying evolutionary processes), it is instrumental to develop theories of spatial decision-making. Here, we present a framework specifically developed to address this issue by the means of artificial neural networks and genetic algorithms. Specifically, we investigate a simple task in which single agents need to learn to explore their square arena without leaving its boundaries. We show that agents evolve by developing increasingly optimal strategies to solve a spatially-embedded learning task while not having an initial arbitrary model of movements. The process allows the agents to learn how to move (i.e. by avoiding the arena walls) in order to make increasingly optimal decisions (improving their exploration of the arena). Ultimately, this framework makes predictions of possibly optimal behavioural strategies for tasks combining learning and movement.

## 1 Introduction

Animals display a diversity of complex behaviours, both social and in relation with their physical environment. Foraging, courtship, predator avoidance, learning and so on are a few examples of the behaviours in which animals engage – deciding at any particular time which activities are suited to their current social and physical contexts.

Natural selection is expected to shape the decision-making processes of animals such that their behaviours are optimised to accomplish fitness relevant goals, within the constraints the animals face. In many circumstances, decisions are made through movements, an animal moving, for instance, towards a food source rather than towards a member of its group. Any constraints altering movements, such as locomotor or cognitive abilities or environmental forces (e.g. water or air flow or obstacles) may therefore be entangled in a decision-making process observed in fine details. Decision-making processes combined with movement involve a great variety of behaviours across taxa, including social insects building behaviour [1], the effect of the environment on animal movements [2], predator detection and avoidance [3], learning experiments in a spatial context [4] and collective motion [5,6]. To evaluate the optimality of decision-making processes when movement is involved is not trivial to formulate mathematically, especially given the high number of factors at play. There has been extensive research on Lévy-walk models to investigate what are the optimal movement patterns to optimise foraging [7–10].

This research effort however faces several limits to understand animal movements at the scale of the individuals and their behaviours in response to biotic and abiotic stimuli [11, 12]. Here we investigate a class of models to explore how rules of interaction emerge from primary goals (finding food, avoiding predators or mating), given a set of constraints

regarding cognitive abilities (in terms of perception or learning) or locomotor capacities. Our aim with this framework is therefore to find out whether or not certain behavioural interactions can arise from first principles considerations.

This framework is based on self-propelled agents imbued with cognitive abilities by means of an artificial neural network which, given a set of stimuli regarding an agent’s environment, determine how the agent moves [13, 14]. During their lives, the fitness of agents is calculated from objective functions representing primary goals and the parameters of this artificial neural network evolve via a simple evolutionary algorithm favouring agents with high fitness. This framework has been used in movement ecology, as a paradigm to study how animals explore and navigate in their environment on relatively long time and large spatial scales [13, 15]. Similar approaches have explored the behavioural rules underlying the emergence of collective behaviours with reinforcement learning [16, 17]. In these studies, the goal of the optimisation process is to find individual-level behavioural rules responsible for specific collective configurations observed in natural flocking systems. For instance, in [17], agents have a cognitive model based on reinforcement learning where the goal is to flock as a rotating ball, a rotating tornado, a rotating hollow core mill, or a rotating full core mill.

However, in animal societies, flocking patterns emerge from behaviours of individual trying to achieve different goals, such as finding food, avoiding predators or mating [18]. In other words, in the existing literature of agent-based models in which agents have a cognitive model, the behavioural rules underlying collective patterns are not *emerging* from first principles (foraging, predation or reproduction) and the reasons *why* animals would behave likewise are still unknown. We note an early attempt of addressing these goals in the literature, which lacked details in the quantification of the emerging behaviours and collective patterns [19]. Investigating the rationale behind the proximate causes of animal spatially-embedded decisions is crucial to get an integrated understanding of animal movements, including collective behaviours, even on small temporal and spatial scales.

In this article, we introduce this class of models for the study of animal behaviour in general and of the cognitive bases on animal movements in particular. To stress our intention to use this class of models for the study of animal behaviour, we name it *smart self-propelled particles*, an explicit reference to self-propelled particles, agent-based models widely used to investigate collective motion [20–22]. Here, we describe the framework in detail, and show how to use it for the study of spatially-embedded animal behaviours. Furthermore, we demonstrate and discuss an approach to develop the models based on empirical data and results. In the present work, we will investigate the optimal movement behaviours in a bounded space, given different fitness functions. We will also investigate the respective effects of different locomotor constraints in affecting the emerging rules of interaction.

## 2 Methods

The framework we introduce for the study of animal behaviour consists of a class of agent-based models, with cognitive agents. “Cognitive agents” refers to the movements of agents being controlled by their brain (an artificial neural network) given a set of stimuli. The parameters of these brains are optimised with a genetic algorithm to achieve a particular task given an ecological context as well as locomotor and cognitive abilities. In this class of models, we expect the structure of the brains (artificial neural networks) to be important in determining the outcome of their predictions. The structure concerns the number of neurons, their arrangement (distributed in one or several so-called hidden layers) and their connections. One possibility to design an architecture of the artificial neural network appropriate to the problem under consideration is to optimise the structure of the network with a genetic algorithm, as part of the brain optimisation [23]. Another simpler approach is to incrementally increase or decrease the number of neurons and to constrain the structure with empirical approach. The latter is the chosen approach in the current study. It allows to focus on the effect of the optimisation of the network parameters in this first investigation but also to closely monitor the effect of structural changes and to potentially infer the cognitive difficulty of a task.

### 2.1 Case study: the emergence of a wall-avoidance behaviour

In this study, we will focus on one particular situation: a single individual moving in an enclosed set-up (e.g. a tank or a cage). Although this seems to be an artificial context, this situation reproduces the context of animals in captivity, enclosed in an arena or a tank, a situation of wide interest in Ethology given the number of experimental designs (for instance involving learning or personality research) set in such environments. Moreover, there are ecological contexts in the wild in which the study of movements when facing boundaries are particularly relevant, such as flying insects avoiding spider webs during their flight. By choosing this situation in this study, we pave the way towards more complex contexts in the future.

Here, cognitive agents are navigating alone (i.e. without interactions with others) in a square arena. They have a life span and their trial either stops (i.e., they die, in genetic algorithm jargon) when hitting the arena walls or at the end of their life time. We explore how different constraints and goals affect the emerging rules of interaction of the agent with the walls and how this relates to empirical results found in fish.

### 2.2 Perception, decisions and movements of agents

#### 2.2.1 Perception

Agents perceive two cues in their environment: their distance from the closest point of any wall and their angle to that wall (Figure 1a). Agents have a simulated brain consisting of an artificial neural network of two input neurons, an hidden layer and one output neuron. The two input neurons correspond to each cue the agents perceive (distance and angle relative to the closest wall), normalised between 0 and 1.

**Figure 1:**
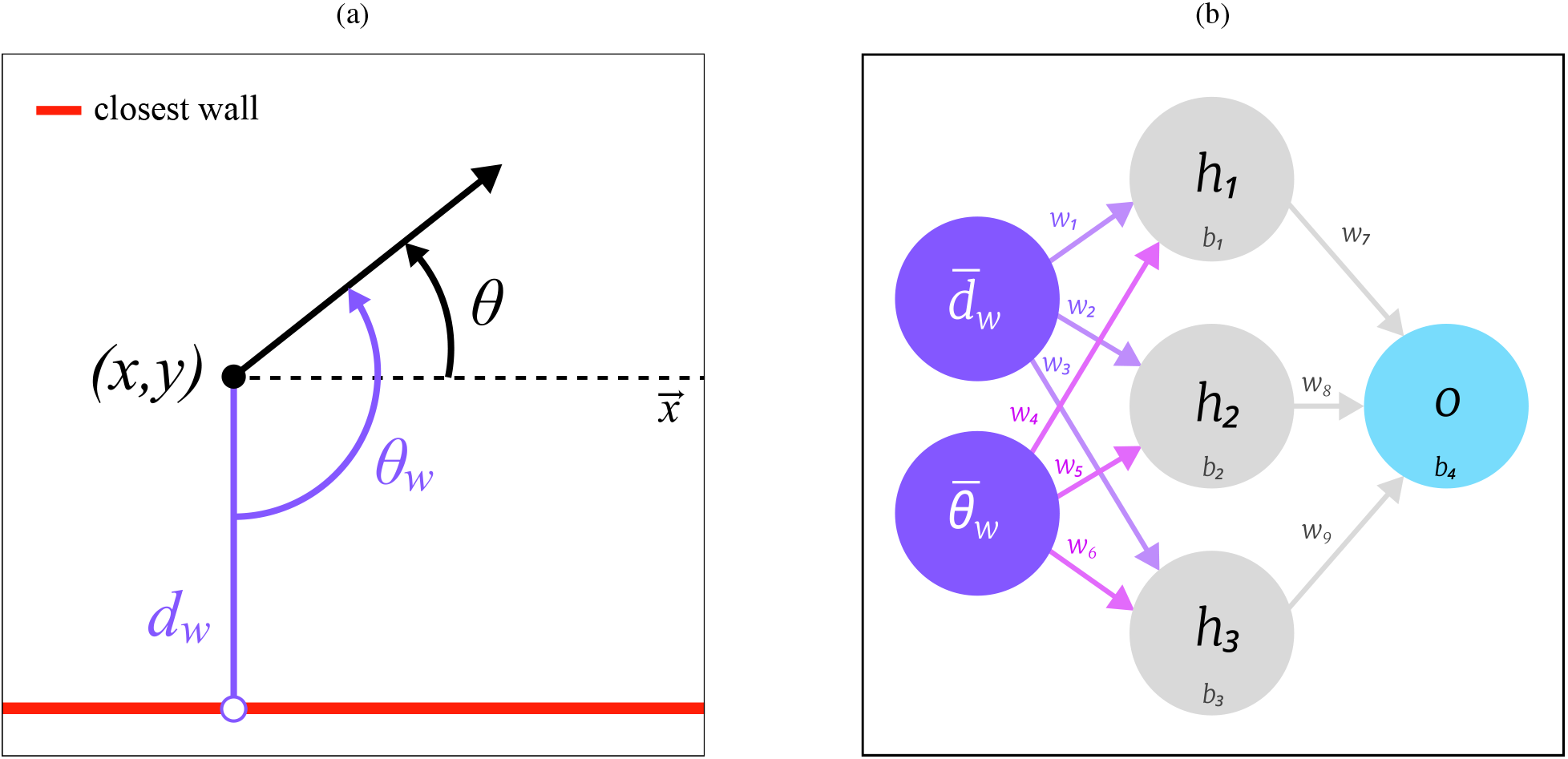
Notations used to depict the location of an agent relative to the closest wall and structure of the artificial neural network. a). The agent (black dot) is located in (*x, y*) with an heading *θ*. Its distance and angle to the closest point of any wall (white dot) are respectively *d_w_* and *θ_w_*. b). Structure of the artificial neural network of reference in our study, with 2 input neurons receiving the normalised distance and angle relative to the closest point of any wall 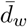 and 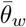, 1 hidden layer with 3 neurons and 1 output neuron. The 13 parameters are made of 9 weights *w_j_* and 4 biases *b_i_*.

#### 2.2.2 Decisions

Decisions regarding movement are made at each time step according to the cues (distance and angle relative to the closest wall) the agent perceives. These cues are transmitted to the input neurons of the artificial neural network of the agent (a multilayer perceptron), which predicts one output value *o*_1_ ∈ [0,1] given by the output neuron. The activation function used in these artificial neural networks is the sigmoid function 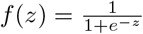, where *z* is the weighted sum of the input connections, added to the bias parameter of the focal neuron (Figure fig:notationsb). The output value of *o*_1_ controls the direction changes of the agent.

The structure of the artificial neural network (i.e. the number of neurons in hidden layers) controls the complexity of the behaviours displayed by the agents. There is therefore a trade-off between the emerging complexity, the generalisation power of the networks and the difficulty to optimise the values of the artificial neural network parameters due to the number of parameters. In this study, we restrict our analysis to networks with one hidden layer. We evaluate, qualitatively, the optimal number of neurons in one hidden layer, between 1 and 6 neurons, by ensuring the chosen number of neurons can at least reproduce the rule of interaction with the wall of actual fish.

#### 2.2.3 Movements

Based on the output value *o*_1_ ∈ [0,1], a coefficient *c*_steering_ = *o*_1_ – 0.5 is calculated (*c*_steering_ ∈ [−0.5,0.5]) and translated into a direction Δ*ϕ* ∈ [−*π, π*]

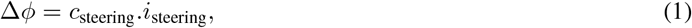

with *i*_steering_ the steering increment *i*_steering_ ∈ [0, 2π], a constant parameter of the model setting the maximal angle turned by agents at each time step. The position (*x, y*) of the agent at time *t* is then updated,

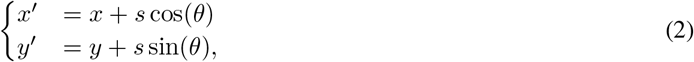

with s the speed of the agent, which is a constant parameter in this study.

### 2.3 Genetic algorithm

#### 2.3.1 Initialisation

At each generation, the same number *n* of individuals is generated. To optimise the exploration of the parameter space across the n agents, each parameter of the artificial neural networks (ie. weights and biases alike) is initialised to a value in 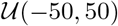 with the Latin Hypercube Sampling method. We use the Python implementation provided by the Surrogate Modeling Toolbox Python package, with the maximin criterion which maximises the minimum distance between points and place the points in a randomised location within their interval [24]. Each agent is simulated four times with different initial locations, randomly taken in the arena, with one initial heading *h* ∈ {0; −*π*; −*π*/2; *π*/2}. A fifth simulation round is executed with the initial location always set to the centre of the arena, and a heading *h* = 0. This additional round is not considered to calculate the fitness of agents but is used to compare the score of agents across generations with a constant initial location, given that the set of initial locations influences the final score in the other four rounds.

#### 2.3.2 Selection

During each round, agents move in a square arena. We have investigated two distinct goals for agents: to survive for the longest time or to maximise the exploration of the arena. With the surviving goal, the score of agents is the sum of time steps during which they managed to stay within the boundaries of the arena. In the exploration goal, agents score one point each time they discover a new cell of the discretised square arena. This mimics a situation where an animal explores an environment in which a resource, such as food, is randomly distributed in space, with a constant probability to be found and collected in any location.

In both goals, the round of an agent ends when the agent goes out of the arena boundaries or if its lifespan exceeds the maximum life span of agents (*T*_max_). When all agents have finished their four rounds, their scores *S* are summed over the rounds and transformed into a fitness.

We investigate the effect of locomotor constraints by focusing on the effects of speed and turning. In many species, turning is costly and affects movement strategies [25–27]. We introduce a penalty to turn, proportional to the squared amount of angle turned in order to penalise very sharp turns. The penalty is set to a constant *p_t_* which is applied at each turn an agent makes such that the total cost to turn across all rounds is *C_t_* ∑ *p_t_* | Δ*ϕ*|^2^. The penalised score *S_p_* = *S* – *C_t_* is then interpolated between 0 and 1, where 0 stands for the minimum score of the generation the current agent belongs to, and 1 for the maximum of the current generation.

The normalised fitness of an agent *i* is

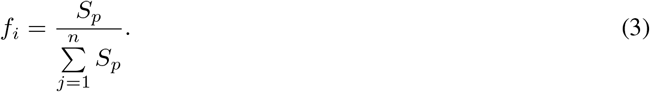

The new generation of n agents is generated from the previous generation by favouring the highest fitnesses. This sampling is stochastic (with replacement), with probability of getting picked proportional to the normalised fitness. Each individual picked to get through the next generation gets the parameters (weights and biases) of its artificial neural network copied and mutated. Small and large mutations occur randomly (33% of the parameters affected by small mutations, 1% by large mutations), by applying a Gaussian noise of standard deviation 0.5 and 5 respectively. These mutation levels have been set by trial and error and the standard deviations have been scaled to the order of magnitude of the parameter values which lie in [−50,50].

### 2.4 Simulations

We simulate the model to investigate its predictions. Each generation consists of *n* = 20000 agents navigating a 600×600 pixels square discretised in a 400-cell grid. Each round can last up to 5000 time steps if the agent does not leave the square boundaries beforehand.

Each simulation runs for 150 generations and is repeated in 60 trials. Parameters explored in our simulations are listed in Table 1. Simulations are implemented in Python 3.10 and analyses performed with both Python and R 4.1.3. We infer rules of interaction with the wall by looking at the average turning angle as a function of the distance and angle to the closest wall. Average turning angles are all calculated using the circular mean

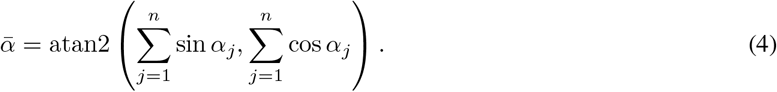

**Table 1:** List of parameters of all simulations. Each row stands for a set of simulation including 60 trials with different initial conditions, each trial having 20000 agents simulated for 150 generations with 5000 time steps each.

| Name | Max. Turning Angle | Nb of neurons in HL | Goal | Speed | Turning penalty |
| --- | --- | --- | --- | --- | --- |
| Parameters of reference | 6.28 | 3 | Explore | 50 | 0 |
| Survive objective function | 6.28 | 3 | Survive | 50 | 0 |
| Parameters with a turning penalty | 6.28 | 3 | Explore | 50 | 0.33 |
| Parameters with a slow speed | 6.28 | 3 | Explore | 5 | 0 |

### 2.5 Empirical data

We use results from a study published in the rummy-nose tetra *Hemigrammus rhodostomus* [28]. This study provided a method to infer rules of interaction with a wall from trajectories of a single fish. We use these results (i) to validate the structure of our artificial neural network, (ii) as an empirical reference to our own simulated results and (iii) use their method to analyse simulated trajectories to gain further understanding on the relationship between behaviour, trajectories and analyses.

## 3 Results

Before analysing simulations in which we varied the goals and constraints of cognition and locomotion, we validated the structure of the artificial neural networks used as artificial brains of the agents in our model. In other words, the capabilities of an artificial neural network structure in predicting empirical results was evaluated, by varying the number of neurons in the hidden layer. Our goal was to have a minimal number of parameters (i.e. a small artificial neural network) to facilitate convergence to optimal solutions and to get a parsimonious and tractable model, while having an artificial brain structure allowing actually observed behaviours to emerge.

### 3.1 Validating the artificial neural network structure

To validate the structure of the artificial neural networks, we checked what would be the least complex artificial neural network that could be trained to predict a known and measured wall-avoidance behaviour of fish swimming in a circular tank [28]. The minimal artificial neural network structure consists of one hidden layer made of one neuron. We trained artificial neural networks with one hidden layer with a number of neurons ranging from 1 to 6 neurons. Each artificial neural network is trained to fit the (simplified) rule of interaction with the wall of *H. rhodostomus*. We visually assessed that a hidden layer made of at least 3 neurons is sufficient to broadly reproduce the behaviour measured in *H. rhodostomus* (Figure 2). Networks with fewer neurons could not completely reproduce the empirically measured behaviour, and networks with more neurons did not qualitatively improved the fit (Electronic Supplementary Material, Figure 1). This validation process is effectively a method to evaluate the cognitive difficulty of any given task. By considering iteratively the effect of the number of neurons, the minimal number of neurons required to achieve a particular task in a specific manner can be inferred and used as an effective measure of the cognitive difficulty of a task. We therefore use this structure (one hidden layer made of three neurons) in the subsequent sections.

**Figure 2:**
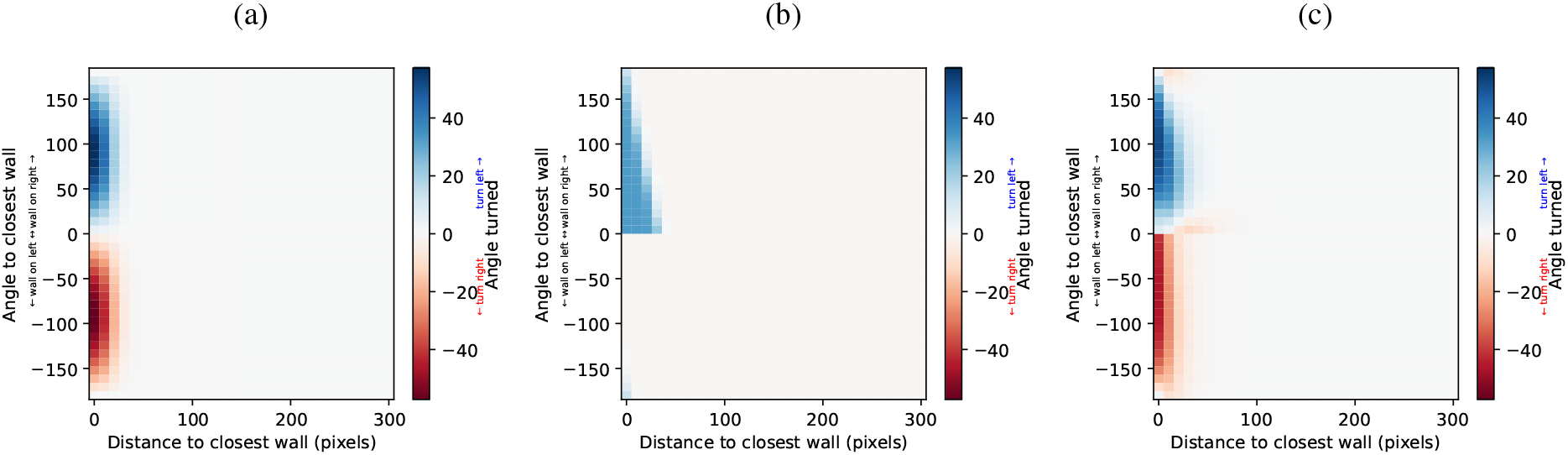
Possibilities of the artificial neural networks used as brains of agents. a). Simplified empirical rule of interaction of fish with the closest point of a circular wall found in *Hemigrammus rhodostomus* [28]. The figure shows how much an agent turns right (red) or left (blue) depending on its position (angle and distance) relative to the closest point of the wall. An angle of 0 represents an agent orthogonally facing the wall. b). Fit of an artificial neural network with one neuron in the hidden layer with the empirical rule of interaction (c). d). Fit of the artificial neural network of reference (Figure 1b) with the empirical rule of interaction (a).

### 3.2 Influence of the objective function

Objective functions define how the fitness of agents is calculated. Fitness is used to favour agents with the highest fitness scores in generating the following generation. To test the effect of the objective function on the emergence of rules of interaction, we defined two different ecological contexts of the goal of a single agent in a bounded environment:

1. the agent has to survive for the longest period of time (i.e. the largest number of time steps);
2. the agent has to explore its environment.

In both situations, as soon as an agent leaves its bounded environment (i.e. its arena), its life stops and so ends its capacity to improve its fitness.

#### 3.2.1 Survive for the longest period of time

In this first objective function, the fitness score of an agent corresponds to the number of time steps in which the agent stays within the boundaries of its arena. The resulting best rules of interaction with the wall averaged across generations and trials (Figure 3a) shows almost no effect of the distance to the closest point of the wall. The trajectories of optimal agents in this situation consists of back-and-forth segments of the shortest length (50 pixels) obtained by doing U-turns at the end of each segment bout (Figure 3b). The directions of these sharp U-turns made to stay close to a segment-like trajectory are on average performed away from the wall (i.e. turning left when the closest wall is on the right-hand side and right when the closest wall is on the left-hand side). Trying to move back-and-forth on a segment is a strategy effectively minimising the explored area for agents moving at a constant speed (that is, not being able to slow down or to stop), therefore minimising the risks of hitting a wall. This is effectively analogous to a freezing behaviour for agents that cannot stop moving. Freezing behaviour is a very common strategy of animals facing predatory threats [29, 30]. Although this population-level strategy differs greatly from the empirical rule of interaction observed in fish, we already see the emergence of the anti-symmetry resulting in agents turning away from the closest wall.

**Figure 3:**
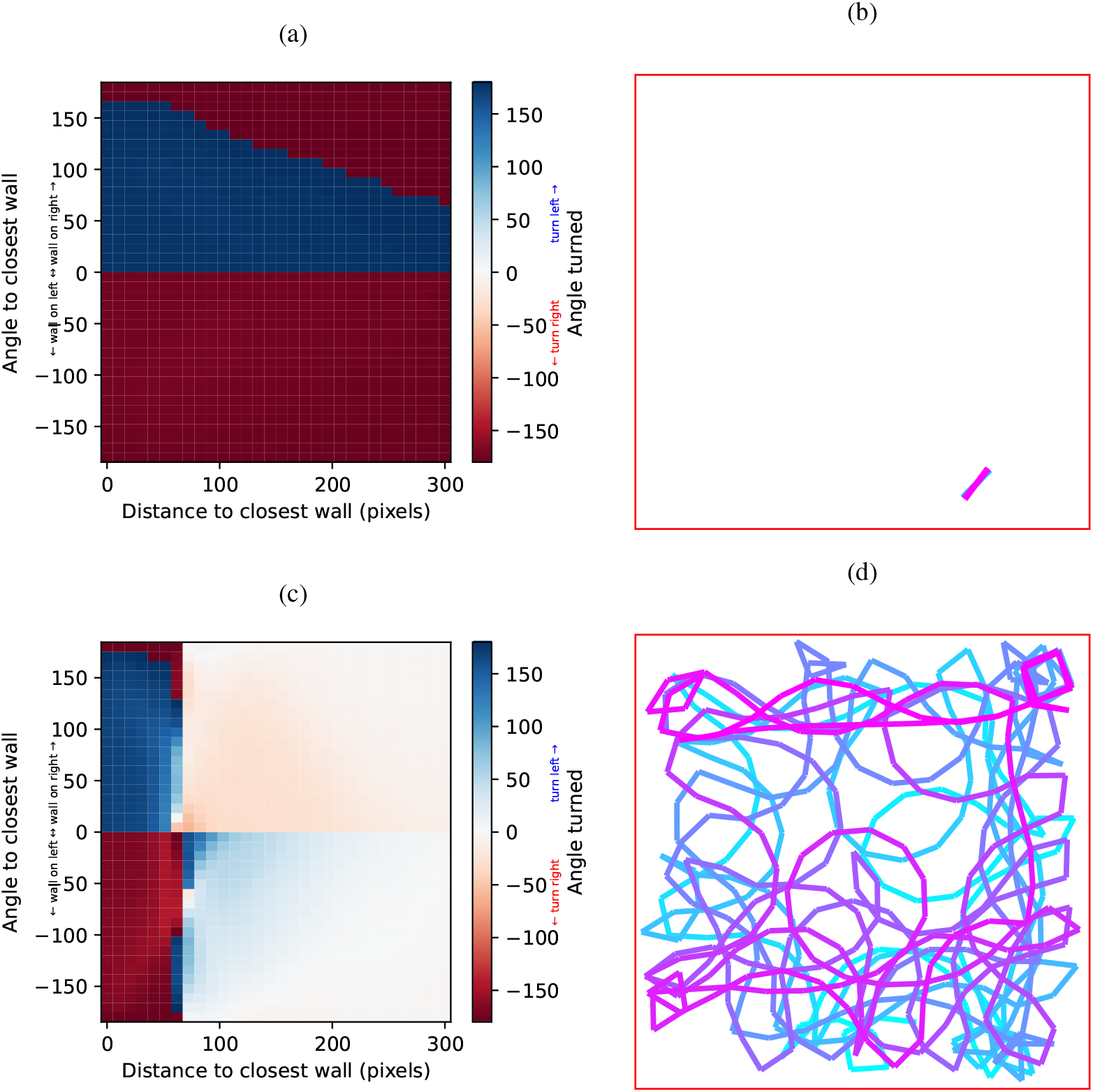
Effect of the objective function on the emergence of the interaction rules. a). Best rules of interaction with the wall averaged across generations and trials when the objective is to survive for the longest period of time. b). Example of a trajectory of an agent trained with the objective to survive for the longest period time. Colour variation depicts the passage of time. c). Average of the best rules of interaction with the wall when the objective is to explore the arena. d). Example of a trajectory of an agent trained when the objective is to explore the arena. Colour variation depicts the passage of time.

#### 3.2.2 Explore the arena

An alternate ecological context is for the agents to increase their fitness when exploring this bounded environment. In this case, the fitness score of an agent corresponds to the number of cells of the discretised arena explored *at least* once (i.e. exploring the same cell several times does not increase the score).

We find that in this context, there is a strong interaction between the angle to the closest point of the wall and the distance to the closest point of the wall (Figure 3c). In agreement with the rules of interaction measured in experiments with *H. rhodostomus*, we find that at the population level, when close to the wall, the direction at which an agent turns (i.e. left or right) depends on its angle to the wall, turning left (respectively right) when the wall is at its right-hand side (respectively left-hand side). This corresponds to a wall-avoidance behaviour, as measured empirically. We also note a weak interaction of attraction to the wall at distances from the wall corresponding to slightly more than the smallest path length of 50 pixels. The optimal trajectories found in this context result in clear wall-following behaviour (Figure 3d), with visits towards the centre of the arena to increase the number of cells of the discretised arena explored. This is an already known result found in fish and bacteria with run-and-tumble motion, in which a wall-avoidance behaviour coupled with discrete reorientations actually results in wall-following behaviour [28, 31, 32]. This alternate objective function produces at the population level interaction rules which are very different from the previous one, in which agents simply needed to survive for as long as they could. This highlights how the definition of the ecological context in our framework affects the resulting optimal rules of interaction.

We now turn to finer quantitative details regarding the movements of agents optimising the exploration criterion. We find that in most of the cases, agents are found at a distance of more than 50 pixels away from the closest wall (Figure 4a). The exception is when agents are facing the closest wall (angle to the wall around 0): in these cases, agents can be found closer to the wall. Three main behaviours regarding angle turned can be identified: very small angle turns, angle turns close to 90 degrees and U-turns (Figure 4b). When agents are close to the wall (within 50 pixels from the wall), they favour turning away from the wall, specifically with large angles, around 90 degrees and, mainly, complete U-turns (Figure 4c). For agents far away from the wall (at least at 150 pixels from the wall), we observe the three modes of the global distribution (small angles, right angles and U-turns), with the predominance of small turning angles (Figure 4d). We note that the distribution of these small angles (less than 60 degrees in magnitude) is compatible with a Normal distribution (Figure 4d inset). This shows the emergence of an apparently stochastic behaviour at the population level – with agents turning as often to the right as to the left when far away from the wall – in a strictly deterministic model of movement. This is in agreement with empirical results found in *H. rhodostomus* [28].

**Figure 4:**
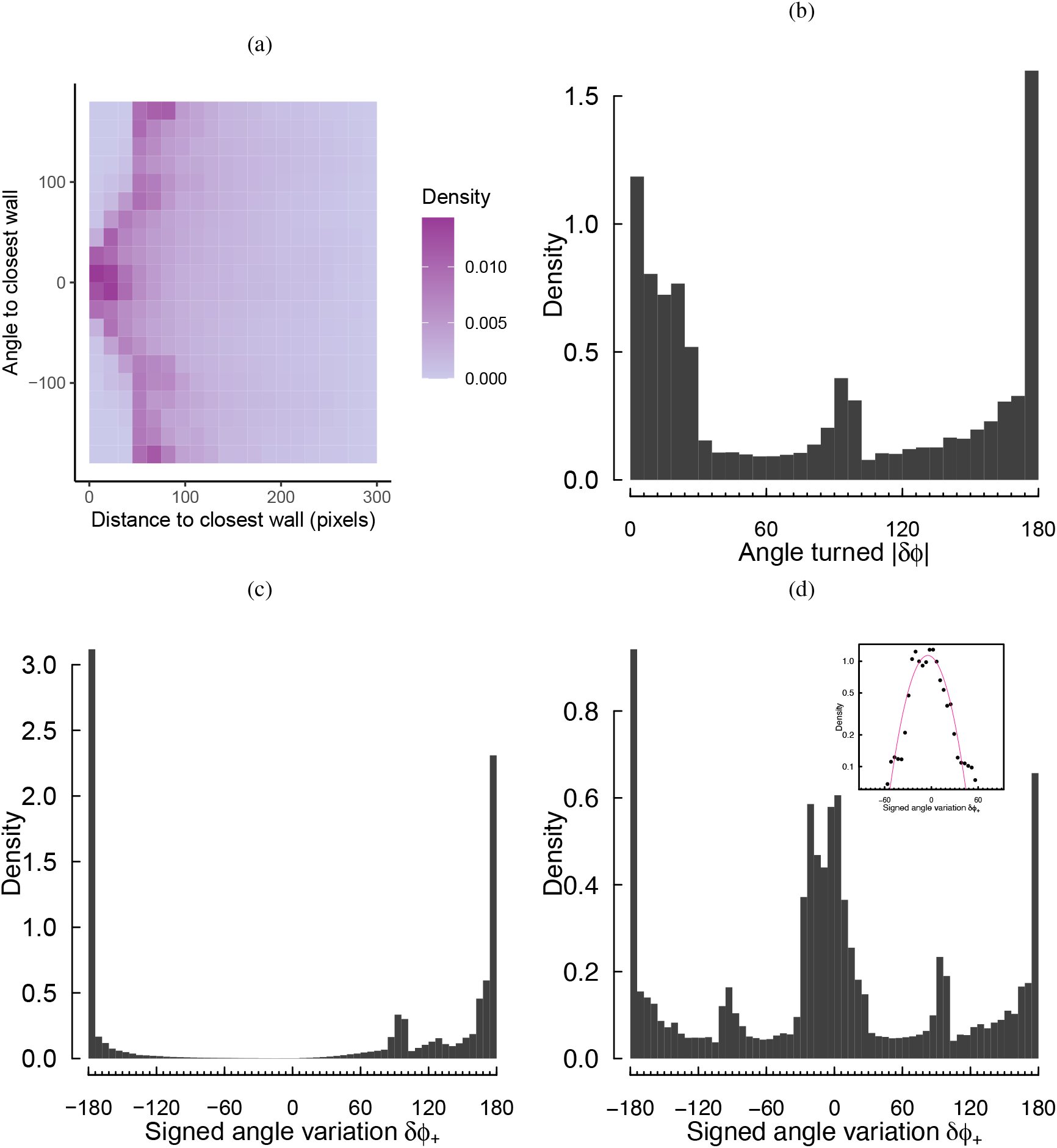
Statistics of exploration and angle changes. a). Density of presence of the best agents as a function of distance and angle to the closest wall. b). Distribution of the absolute value of the angle turned. c). Distribution of the signed angle variation restricted to agents close to the wall (i.e. less than 50 pixels from the wall), *δϕ*_+_ = *δϕ* × sign(*θ_w_*), positive for angle turned away from the wall, negative otherwise. d). Distribution of the signed angle variation restricted to agents far away from the wall (i.e. more than 150 pixels from the wall). Inset: zoom of the distribution between −60 and 60 degrees. Purple line shows a Gaussian fit.

### 3.3 Individual-level interaction rules vs population-level interaction rules

The average behaviour shown Figure 3c is not the one actually used by the agents (Figure 5). We can indeed find equally good performances by individuals whose rule of interaction with the wall greatly differ from the average one. The behaviour of these agents is very similar in regions of the input variable space under strong selection – here, when agents are close to a wall, we see a strong wall-avoidance behaviour. However, when the agents are far from a wall and therefore in regions under less selection pressures, we see more variation. For instance, the distance at which individuals start to avoid the wall or the strength or existence of a parameter region in which agents are attracted to the closest wall. This leads to a diversity of behaviours and forms of trajectories even though, on average, agents all successfully avoid the wall by turning left when the wall is on their right and vice versa. This suggests the possibility for diversity in the presence of strong selective pressures, where early experience leads to a diverse yet equally satisfying pool of solutions. It is worth noting that this diversity is difficult to notice in empirical data and generally overlooked. Most of the empirical studies investigating rules of interaction in animals report results obtained from behaviours averaged across individuals, trials, time of the day and days of experiment [33–36], preventing the finding of diverse solutions from different individuals or depending on statuses hidden to the experimenter, such as the state of hunger of a specific individual.

**Figure 5:**
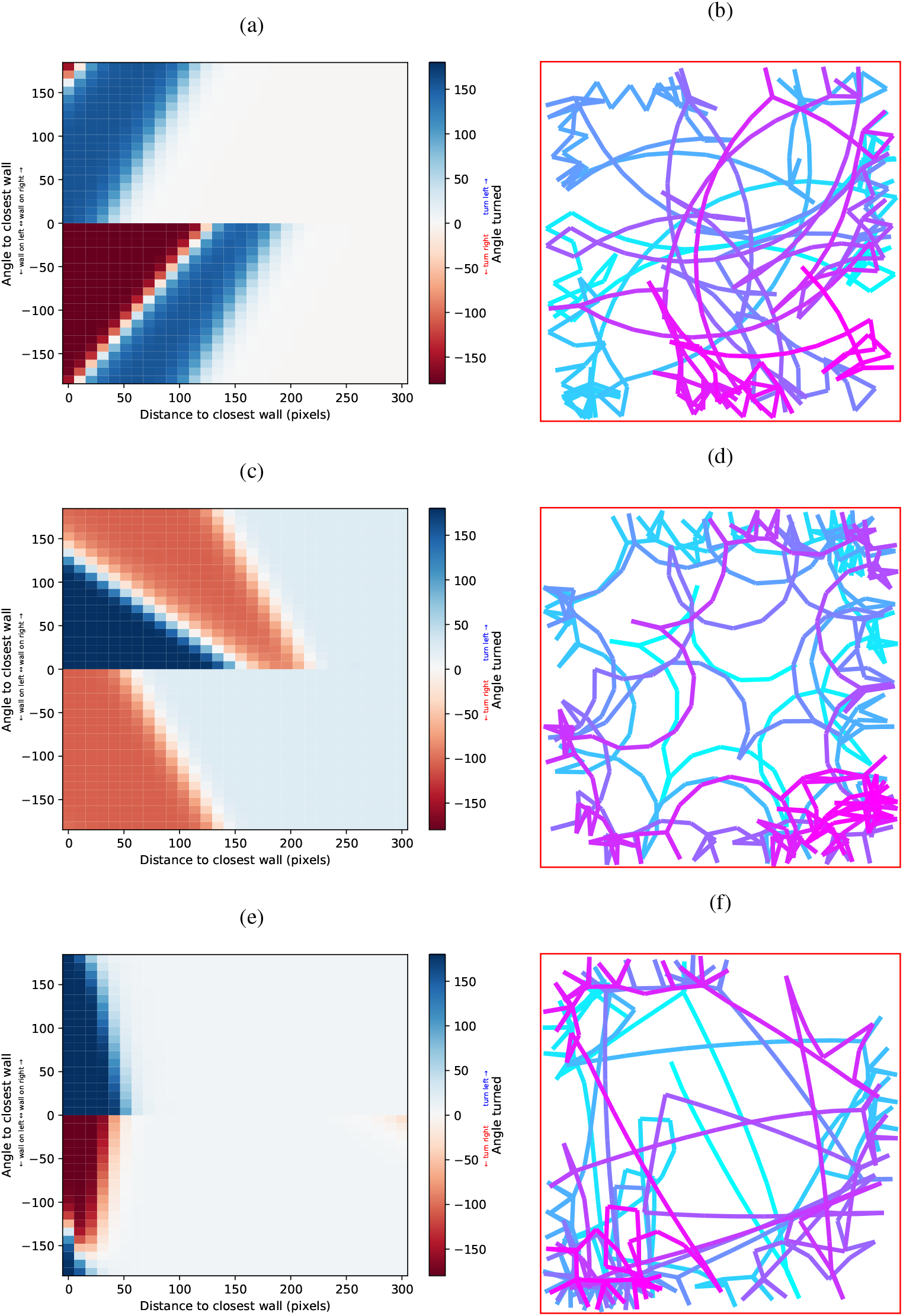
Diversity of equally efficient strategies to solve the exploration problem. a, c and e). Angle turned as a function of the angle and distance to the closest point on the wall of three distinct agents, solving the exploration problem with the maximum exploration score. b, d and f). Trajectories corresponding to, respectively, rules of interaction (a), (c) and (e). Colour variation depicts the passage of time.

### 3.4 Effect of locomotor constraints

We study the effect of locomotor constraints by investigating the costs associated with turning (Figure 6a-c). We find that overall, the optimal behaviour stays similar: agents still need to turn left when the wall is at their right hand-side and vice versa (Figure 6a). We note that agents start turning earlier (i.e. further away from the wall) when they get to face the wall and turn later on (i.e. at closest distances from the wall) when they do not directly face the wall, unlike the rule of interaction of reference (Figure 3c). This is likely the result of agents minimising the instances when they need to turn a lot to avoid the wall.

**Figure 6:**
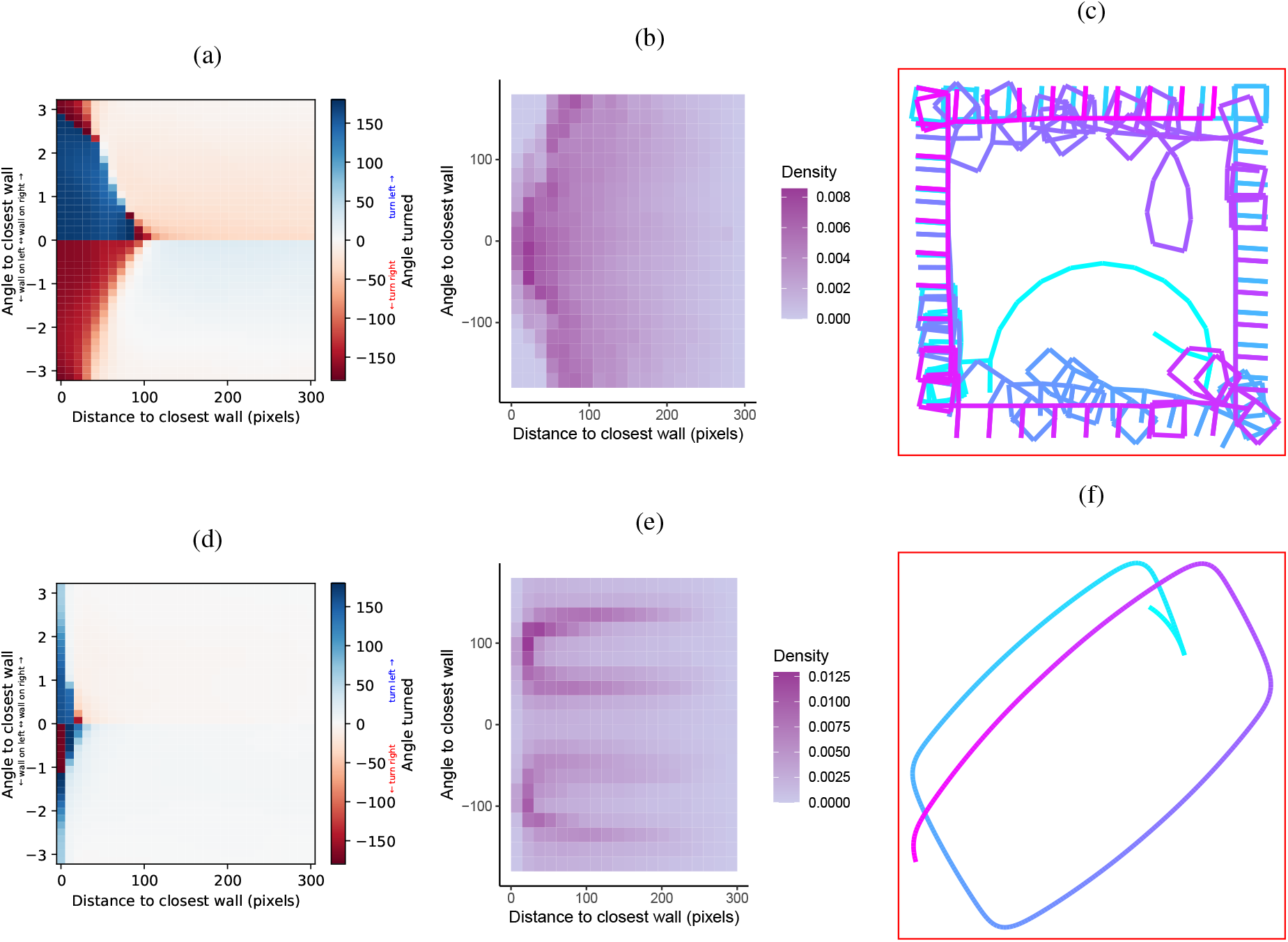
Effect of locomotor constraints on the optimal rule of interaction with the wall. a-c). Results when introducing a turning penalty every time the agent turns. d-f). Results when agents are moving 10 times slower than the speed of reference (5 pixels per time step in contrast to 50). a and d). Best rules of interaction with the wall averaged across generations and trials. b and e). Density of presence of the best agents as a function of distance and angle to the closest wall. c and f). Example of trajectories of the best agents of each locomotor constraint. Colour depicts the passage of time.

We also investigated the effect of speed by running simulations in which agents move ten times slower than in the conditions of reference – namely 5 pixels per time unit instead of 50 (Figure 6d-e). This change in the locomotor capacities relative to the size of the bounded space affects mainly the distance at which agents start to avoid the closest wall (Figure 6d). Agents also make much smaller turns, resulting in a distribution of angle turned very different from the distribution obtained with the parameters of reference (ESM, Figure 4a). This results in trajectories being less tortuous than other rules of interaction found for previous constraints (Figure 6f). This leads agents to being distributed closer to the wall on average (ESM, Figure 4b) with two clear angles relative to the wall around 54 and 132 degrees.

## 4 Discussion

We find that the objective function affects the movement behaviours that emerge, with a great variety of solutions found to the same problem. In particular, we show that a wall-following trajectory emerges from wall avoidance behaviour of agents actually optimising the exploration of their bounded environment. They do so by successfully learning how to avoid the obstacles of their environment, based on very simple local cues. The emerging rules of interaction with the wall resembles the one found in the fish *H. rhodostomus*, where single fish were studied when swimming freely in small circular tanks [28]. At the population level, we find indeed a clear anti-symmetric wall avoidance behaviour at close distances from the wall, in agreement with empirical data [28]. This is achieved with an extremely simple cognitive model made of an artificial neural network with only 13 parameters, and whose input is also kept very simple – only information regarding the position of the agent relative to the closest point of any wall of the arena is provided. Locomotor abilities such as the speed or turning abilities also affect the emerging behaviours, changing both the interplay between distance and angle to the wall in determining wall avoidance and the resulting trajectories and arena exploration.

We stress the importance of combining the use of this class of models with empirical data. Such a theoretical approach provides multiple strategies, many of them potentially being biologically unrealistic. Empirical data therefore provide additional context and constraints to guide the development of meaningful models. Moreover, adding constraints reduces the diversity of optimal strategies successfully solving the problem under study and helps excluding sub-optimal solutions.

The good agreement between our findings and empirical results in fish occur at the population level, on aggregated and averaged simulated data. However, this fit overlies a complexity largely unexplored in biology. Our simulation results look similar once averaged across multiple agents but the individual behaviours actually encompass an important variety of strategies for exploring the arena while avoiding the wall, although not in high-risk zones, which are less diverse in the exhibited behaviours. The discrepancy between individual and population level behaviours and the heterogeneity of behaviours at the individual level are rarely investigated when looking for rules of interaction while it could occur that none of the individual actually behaves as the average behaviour. Identifying whether the average behaviour is exhibited by many individuals may allow to identify behaviours emerging from either high or low selective pressures. Our results obtained with an objective function requiring individuals to maximise their time spent without touching the wall suggest that freezing behaviours are compatible with individuals trying to favour survival against exploration when facing extreme threats [29, 30].

In our study, we find apparently stochastic behaviour emerging at the population level: when agents are moving far away from the closest wall, their average behaviour is Gaussian and apparently stochastic at the population level – they turn as often to the left as to the right on average. This apparently stochastic behaviour emerges in a model of movement which is strictly deterministic: when submitted with the same cues, the agents will always move exactly in the same way. Stochastic behaviours are often interpreted as the signature of free will or noise in empirical data [28]. Our result here stresses that, in practice, one needs to be careful when pooling empirical data from multiple individuals together, e.g. to calculate an average behaviour: apparent stochasticity may arise even when agents with various strategies have a rationale and unnoisy behaviour. This underlines the importance of providing evidence that animals exhibit a stochastic behaviour at the individual level rather than at the group or population level.

Our results highlight the value of our approach to test the direct effect of an ecological context in evaluating the optimality of rules of interaction of animals with their social and physical environments. It is an holistic approach providing a class of models to explore how animals solve tasks, fulfil their needs, in the presence of complex trade-offs and constraints brought by their environment and their own perceptory, cognitive, physiological and locomotor machineries. This class of models therefore fills a gap by providing a method to investigate systems combining two distinct aspects of spatially-embedded behaviours: decision-making and movements. In other words, this class of models allows integrated research of the cognitive bases of animal movements, specifically by unifying two research bodies often divided. For instance, in the field of collective motion which studies how groups of animals such as bird flocks or fish school coordinate and move in a synchronised manner, the literature is split between studies focusing on the decision-making underlying movements in groups and studies looking at the spatial mechanisms of movements. Here, we propose to explore optimal decision-making *under the constraints* of spatially-embedded tasks – that is taking into account the respective effects of intent, perception, locomotion and obstacles. Although this class of models preexisted our study [13], it was mainly used to study the optimality of animal movements on large spatial and temporal scales, i.e. on scales on which the resolution of empirical data does not allow the study of cognition – or lacked analysis of emerging behaviours in relation to empirical data [14, 19].

We show here how this framework can be used with empirical data on fine temporal and spatial scales, to both constrain the exploration and guide the modelling choices to successfully study the cognitive bases of animal movements. The smart self-propelled particles approach potentially benefits several scales of description. In addition to its use in Behavioural Ecology to investigate animal movements, for instance investigating the optimal spatial distribution of animals given a set of constraints, the class of models can also be applied in Ethology, for instance to compare experimental results of learning experiments involving movements with theoretical predictions. While most empirical studies are descriptive and do not really explain why animals behave the way they do, we propose here, from the descriptive empirical knowledge of animal behaviours, to reverse engineer the fitness functions shaping the behaviours of animals. Doing so will help understand why animals adopt the rules of interaction predicted by the smart self-propelled particles framework. On the other hand, should the predictions differ from empirical observations, it will help formulating improved model by adding or removing constraints or changing the goal agents have to achieve, essentially inferring the fitness function of the animal. This study paves the way towards the development of normative models aimed at understanding better the cognitive bases of decision-making of moving animals with an holistic and integrated approach allowing to evaluate the respective effects of cognitive, locomotor and environmental constraints.

## Supporting information

Electronic Supplementary Material

## References

1 A. Khuong, J. Gautrais, A. Perna, C. Sbaï, M. Combe, P. Kuntz, C. Jost, and G. Theraulaz, “Stigmergic construction and topochemical information shape ant nest architecture”, Proceedings of the National Academy of Sciences 113, Publisher: Proceedings of the National Academy of Sciences, 1303–1308 (2016).

2 A. Khuong, V. Lecheval, R. Fournier, S. Blanco, S. Weitz, J.-J. Bezian, and J. Gautrais, “How do ants make sense of gravity? a boltzmann walker analysis of lasius niger trajectories on various inclines”, PLOS ONE 8, Publisher: Public Library of Science, e76531 (2013).

3 D. Rodríguez-Morales, H. Tapia-McClung, L. E. Robledo-Ospina, and D. Rao, “Colour and motion affect a dune wasp’s ability to detect its cryptic spider predators”, Scientific Reports 11, 15442 (2021).

4 M. H. Pillot, J. Gautrais, J. Gouello, P. Michelena, A. sibbald, and R. Bon, “Moving together: incidental leaders and naïve followers”, Behavioural Processes 83, 235–241 (2010).

5 A. J. W. Ward, D. J. T. Sumpter, I. D. Couzin, P. J. B. Hart, and J. Krause, “Quorum decision-making facilitates information transfer in fish shoals”, Proceedings of the National Academy of Sciences 105, 6948–6953 (2008).

6 H. Ling, G. E. Mclvor, K. van der Vaart, R. T. Vaughan, A. Thornton, and N. T. Ouellette, “Costs and benefits of social relationships in the collective motion of bird flocks”, Nature Ecology & Evolution 3, 943–948 (2019).

7 G. M. Viswanathan, S. V. Buldyrev, S. Havlin, M. Da Luz, E. Raposo, and H. E. Stanley, “Optimizing the success of random searches”, Nature 401, 911–914 (1999).

8 G. M. Viswanathan, M. G. Da Luz, E. P. Raposo, and H. E. Stanley, The physics of foraging: an introduction to random searches and biological encounters (Cambridge University Press, 2011).

9 A. James, M. J. Plank, and A. M. Edwards, “Assessing lévy walks as models of animal foraging”, Journal of The Royal Society Interface 8, 1233–1247 (2011).

10 V. Zaburdaev, S. Denisov, and J. Klafter, “Lévy walks”, Rev. Mod. Phys. 87, 483–530 (2015).

11 G. H. Pyke, “Understanding movements of organisms: it’s time to abandon the lévy foraging hypothesis”, Methods in Ecology and Evolution 6, Publisher: John Wiley & Sons, Ltd, 1–16 (2015).

12 A. Reynolds, “Liberating lévy walk research from the shackles of optimal foraging”, Physics of Life Reviews 14, 59–83 (2015).

13 T. Mueller, W. F. Fagan, and V. Grimm, “Integrating individual search and navigation behaviors in mechanistic movement models”, Theoretical Ecology 4, 341–355 (2011).

14 N. Bredeche and N. Fontbonne, “Social learning in swarm robotics”, Philosophical Transactions of the Royal Society B: Biological Sciences 377, Publisher: Royal Society, 20200309 (2022).

15 R. Nathan, W. M. Getz, E. Revilla, M. Holyoak, R. Kadmon, D. Saltz, and P. E. Smouse, “A movement ecology paradigm for unifying organismal movement research”, Proceedings of the National Academy of Sciences 105, 19052–19059 (2008).

16 M. Durve, F. Peruani, and A. Celani, “Learning to flock through reinforcement”, Phys. Rev. E 102, Publisher: American Physical Society, 012601 (2020).

17 T. Costa, A. Laan, F. J. H. Heras, and G. G. de Polavieja, “Automated discovery of local rules for desired collective-level behavior through reinforcement learning”, Frontiers in Physics 8 (2020).

18 D. J. Sumpter, “Collective animal behavior”, in Collective animal behavior (Princeton University Press, 2010).

19 C. R. Ward, F. Gobet, and G. Kendall, “Evolving collective behavior in an artificial ecology”, Artificial life 7, 191–209(2001).

20 I. Aoki, “A simulation study on the schooling mechanism in fish”, Bulletin of the Japanese Society of Scientific Fisheries 48, 1081–1088 (1982).

21 T. Vicsek, A. Czirók, E. Ben-Jacob, I. Cohen, and O. Shochet, “Novel type of phase transition in a system of self-driven particles”, Phys. Rev. Lett. 75, 1226–1229 (1995).

22 U. Lopez, J. Gautrais, I. D. Couzin, and G. Theraulaz, “From behavioural analyses to models of collective motion in fish schools”, Interface Focus 2, 693–707 (2012).

23 F. Leung, H. Lam, S. Ling, and P. Tam, “Tuning of the structure and parameters of a neural network using an improved genetic algorithm”, IEEE Transactions on Neural Networks 14, 79–88 (2003).

24 M. A. Bouhlel, J. T. Hwang, N. Bartoli, R. Lafage, J. Morlier, and J. R. R. A. Martins, “A python surrogate modeling framework with derivatives”, Advances in Engineering Software, 102662 (2019).

25 R. P. Wilson, I. W. Griffiths, P. A. Legg, M. I. Friswell, O. R. Bidder, L. G. Halsey, S. A. Lambertucci, and E. L. C. Shepard, “Turn costs change the value of animal search paths”, Ecology Letters 16, Publisher: John Wiley & Sons,Ltd, 1145–1150 (2013).

26 R. Munden, L. Börger, R. P. Wilson, J. Redcliffe, R. Brown, M. Garel, and J. R. Potts, “Why did the animal turn? time-varying step selection analysis for inference between observed turning-points in high frequency data”, Methods in Ecology and Evolution 12, Publisher: John Wiley & Sons, Ltd, 921–932 (2021).

27 N. J. Klappstein, J. R. Potts, T. Michelot, L. Börger, N. W. Pilfold, M. A. Lewis, and A. E. Derocher, “Energy-based step selection analysis: modelling the energetic drivers of animal movement and habitat use”, Journal of Animal Ecology 91, Publisher: John Wiley & Sons, Ltd, 946–957 (2022).

28 D. S. Calovi, A. Litchinko, V. Lecheval, U. Lopez, A. Pérez Escudero, H. Chaté, C. Sire, and G. Theraulaz, “Disentangling and modeling interactions in fish with burst-and-coast swimming reveal distinct alignment and attraction behaviors”, PLOS Computational Biology 14, 1–28 (2018).

29 D. Eilam, “Die hard: a blend of freezing and fleeing as a dynamic defense—implications for the control of defensive behavior”, Neuroscience & Biobehavioral Reviews 29, Defensive Behavior, 1181–1191 (2005).

30 K. Roelofs, “Freeze for action: neurobiological mechanisms in animal and human freezing”, Philosophical Transactions of the Royal Society B: Biological Sciences 372, 20160206 (2017).

31 I. D. Vladescu, E. J. Marsden, J. Schwarz-Linek, V. A. Martinez, J. Arlt, A. N. Morozov, D. Marenduzzo, M. E. Cates, and W. C. K. Poon, “Filling an emulsion drop with motile bacteria”, Phys. Rev. Lett. 113, 268101 (2014).

32 J. Elgeti and G. Gompper, “Run-and-tumble dynamics of self-propelled particles in confinement”, Europhysics Letters 109, 58003 (2015).

33 J. E. Herbert-Read, A. Perna, R. P. Mann, T. M. Schaerf, D. J. T. Sumpter, and A. J. W. Ward, “Inferring the rules of interaction of shoaling fish”, Proceedings of the National Academy of Sciences 108, 18726–18731 (2011).

34 Y. Katz, K. Tunstrøm, C. C. Ioannou, C. Huepe, and I. D. Couzin, “Inferring the structure and dynamics of interactions in schooling fish”, Proceedings of the National Academy of Sciences 108, 18720–18725 (2011).

35 A. K. Zienkiewicz, F. Ladu, D. A. Barton, M. Porfiri, and M. D. Bernardo, “Data-driven modelling of social forces and collective behaviour in zebrafish”, Journal of Theoretical Biology 443, 39–51 (2018).

36 R. Escobedo, V. Lecheval, V. Papaspyros, F. Bonnet, F. Mondada, C. Sire, and G. Theraulaz, “A data-driven method for reconstructing and modelling social interactions in moving animal groups”, Philosophical Transactions of the Royal Society B: Biological Sciences 375, 20190380 (2020).

