## Supplementary material for "Smart self-propelled particles: a framework to investigate the cognitive bases of movement": Electronic Supplementary Material

**Valentin Lecheval**
School of Mathematics
University of Leeds, UK


**Richard P. Mann**
School of Mathematics
University of Leeds, UK

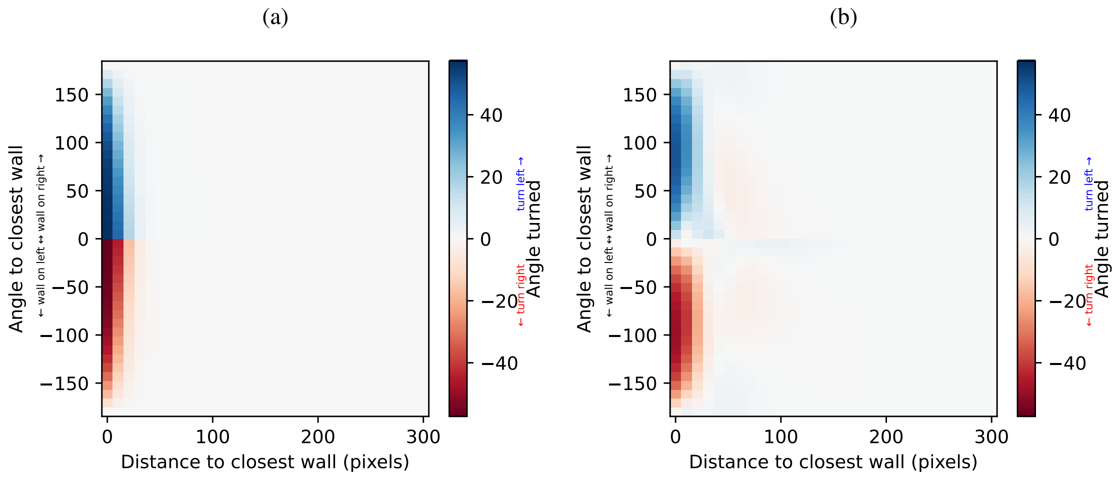


Figure 1: Structure and possibilities of the artificial neural networks used as brains of agents. a). Fit of an artificial neural network with two neurons in the hidden layer with the empirical rule of interaction. b). Fit of an artificial neural network with six neurons in the hidden layer with the empirical rule of interaction

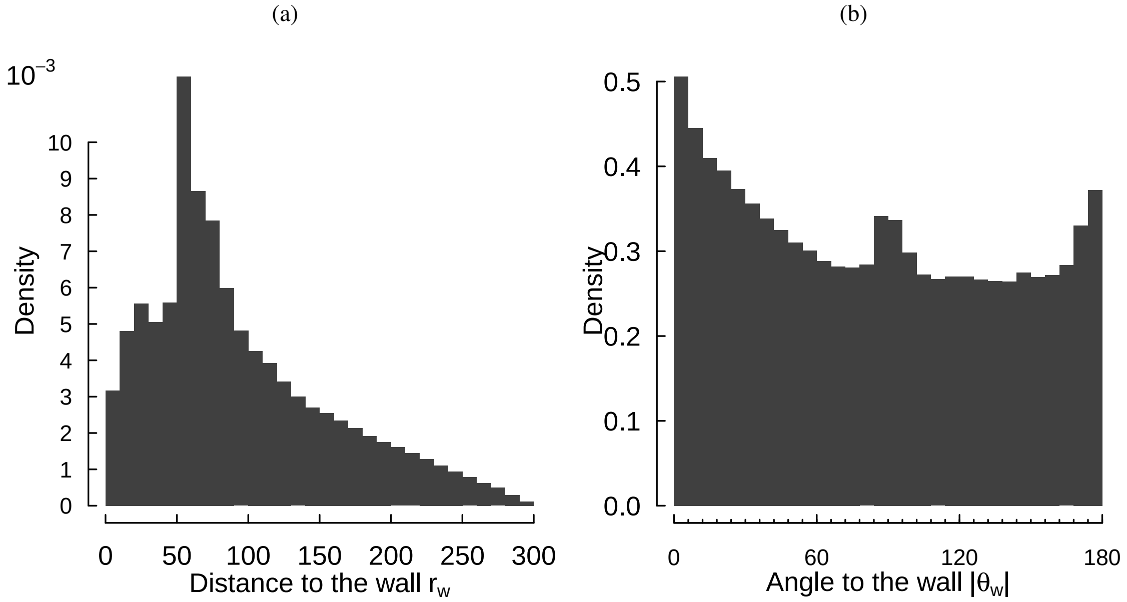


Figure 2: Distributions of the distance (a) and angle (b) to the closest point of the closest wall for best agents from simulations with parameters of reference. Distributions from trajectories of 118 agents resulting in 5384408 data points.

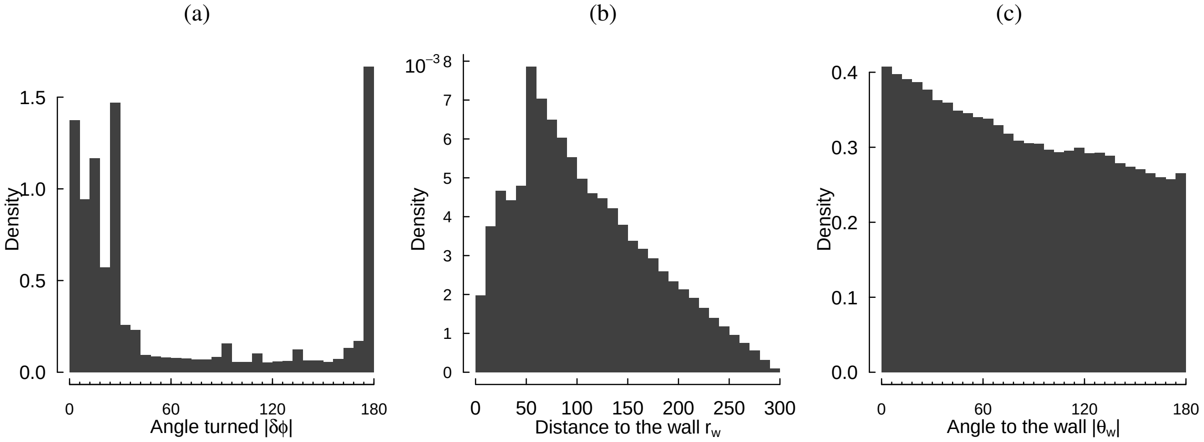


Figure 3: Distributions of the angle turned (a) and distance (b) and angle (c) to the closest point of the closest wall for best agents from simulations with a turning penalty. Distributions from trajectories of 64 agents resulting in 2742239 data points.

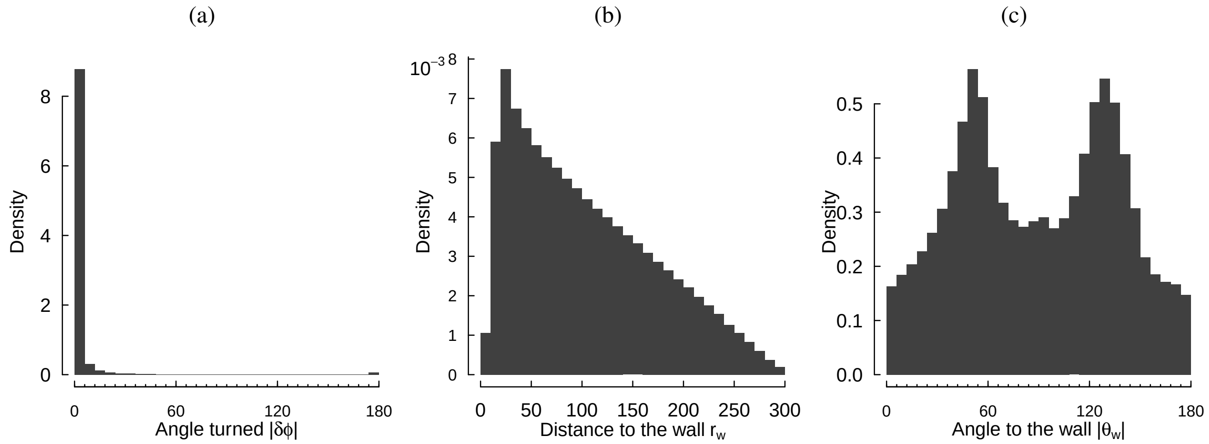


Figure 4: Distributions of the angle turned (a) and distance (b) and angle (c) to the closest point of the closest wall for best agents from simulations with a speed 10 times slower than the speed of reference. Distributions from trajectories of 65 agents resulting in 3239184 data points.
